# Machine Learning Approaches for Predicting Virus-Human Protein-Protein Interactions: An Evaluation of Retroviral Interaction Networks

**DOI:** 10.1101/2024.11.13.623326

**Authors:** Omid Mahmoudi, Somayye Taghvaei, Shirin Salehi, Soheil Khosravi, Alireza Sazgar, Sara Zareei

## Abstract

Virus-human protein-protein interactions (VHPPI) are key to understanding how viruses manipulate host cellular functions. This study constructed a retroviral-human PPI network by integrating multiple public databases, resulting in 1,387 interactions between 29 retroviral and 1,026 human genes. Using minimal sequence similarity, we generated a pseudo-negative dataset for model reliability. Five machine learning models—Logistic Regression (LR), Support Vector Machine (SVM), Naive Bayes (NB), Decision Tree (DT), and Random Forest (RF)—were evaluated using accuracy, sensitivity, specificity, PPV, and NPV. LR and KNN models demonstrated the strongest predictive performance, with sensitivities up to 77% and specificities of 52%. Feature importance analysis identified GC content and semantic similarity as influential predictors. Models trained on selected features showed enhanced accuracy with reduced complexity. Our approach highlights the potential of computational models for VHPPI predictions, offering valuable insights into viral-host interaction networks and guiding therapeutic target identification.

**Significance:** This study addresses a crucial gap in antiviral research by focusing on the prediction of virus-host protein-protein interactions (VHPPI) for retroviruses, which are linked to serious diseases, including certain cancers and autoimmune disorders. By leveraging machine learning models, we identified essential host-pathogen interactions that underlie retroviral survival and pathogenesis. These models were optimized to predict interactions accurately, offering valuable insights into the complex mechanisms that retroviruses use to manipulate host cellular processes. Our approach highlights key host and viral proteins, such as ENV_HV1H2 and CD4, that play pivotal roles in retroviral infection and persistence. Targeting these specific interactions can potentially disrupt the viral lifecycle while minimizing toxicity to human cells. This study thus opens avenues for the development of selective therapeutic strategies, contributing to more effective and targeted antiviral interventions with fewer side effects, marking a significant step forward in computational virology and drug discovery.

## Introduction

The *Retroviridae* family encompasses a wide range of single-stranded RNA viruses (ssRNA) that hold significance in human health due to their roles in disease development. In particular, human endogenous retroviruses (HERVs), have shown associations with several cancers and autoimmune diseases. for instance, HERV elements like HERV-K are expressed in various tumors including breast cancer and melanoma [1]. Additionally, the reactivation of HERV play a role in autoimmune disease such as lupus and multiple sclerosis. Retroviral proteins produced by HERV activation can act as superantigens which overstimulate the immune responses in conditions such as rheumatoid arthritis [2, 3].

Retroviruses interact with host proteins in a variety of ways enabling them to invade, replicate, and persist within the host cells. These interactions guarantee the virus’ lifecycle and also involves the manipulation of cellular structures, pathways, and proteins which is the reason for the its pathogenesis. Retroviral proteins enable the virus entry by binding to specific receptors on the host cells surface. For instance, HIV-1 envelope glycoproteins binds to the host CD4 and CXCR4 receptors [4]. Inside the host cell, retroviruses need to hijack the dense network of cytoskeleton to reach the nucleus which includes the interaction with host proteins [5, 6]. The integration of retrovirus genome into the host genome also needs the interaction of viral proteins with the host one such as karyopherins, which aid in transporting the viral DNA across the nuclear membrane into the host cell’s nucleus [7]. Retroviral integrase interact with (bromodomain and extra-terminal domain) proteins and increase the efficacy of viral genome integration [8]. During genome integration, retroviruses may exploit the host’s DNA repair proteins [9] and interact with the host translational machinery. For example, specific untranslated regions (UTRs) in some retroviruses interact with host RNA-binding proteins, ensuring that the viral protein synthesis is prioritized over cellular protein production [10].

Since viral proteins rely on unique and essential interaction with human proteins to replicate and survive, identifying the protein-protein interaction (PPI) network between host and viral proteins known as the host-virus protein-protein interaction or VHPPI is vital for identifying new drug targets. Blocking these interactions can disrupt the viral pathogenesis without significant negative impact on normal cellular processes, making the anti-retroviral treatments more effective and less toxic [11].

The identification of VHPPI can be done using wet lab techniques, including Förster resonance energy transfer (FRET) [12], Yeast two hybrid screening (Y2H) [13], surface plasmon resonance (SPR) [14], and protease assay [15].

Bioinformatics is the field that combines biology, computer science, and mathematics to analyze and interpret biological data. Its applications include genomics [16-21], proteomics [22-27], and metagenomics [28-31]. It also applies machine learning techniques to build predictive models in personalized medicine [32] or to predict PPIs [33]. In the present study, we developed and optimized machine learning models to predict the HVIN for retrovirus family and evaluated the performance of each model.

## Computational Approaches

### Data Collection

#### Virus-Human Protein-Protein Interaction (PPI) Network

To construct the virus-human protein-protein interaction (PPI) network, we integrated data from multiple public databases including STRING [34] Intact [35], DIP [36], VirusMINT [37], and BioGRID [38] retrieving the retroviral interactions. This integration focused on retrieving interactions involving retroviruses. The resulting dataset was considered as positive interactions which included 29 retroviral genes and 1,026 human genes, generating an interaction network composed of 1,387 interactions.

To construct the negative dataset, we employed a strategy previously applied in studies of other virus-host interactions [39]. Specifically, pseudo-negative samples were generated by selecting pairs of proteins with minimal sequence similarity to those in the positive interaction set using CD-HIT [40] with the threshold of less than 20% sequence similarity: any virus-human protein pairs that shared less than 20% similarity with known positive interaction pairs were considered as candidates for the negative dataset. This approach aims to reduce the likelihood of including true interactions in the negative set, thereby enhancing the reliability of the negative samples.

Finally, to prepare data for model training and evaluation, we split both positive and negative datasets into training and test sets, following an 80/20 ratio.

### Feature Extraction

In this study, we used a range of protein and nucleotide sequence-based features for prediction which are described in Table 1.

**Table 1.**
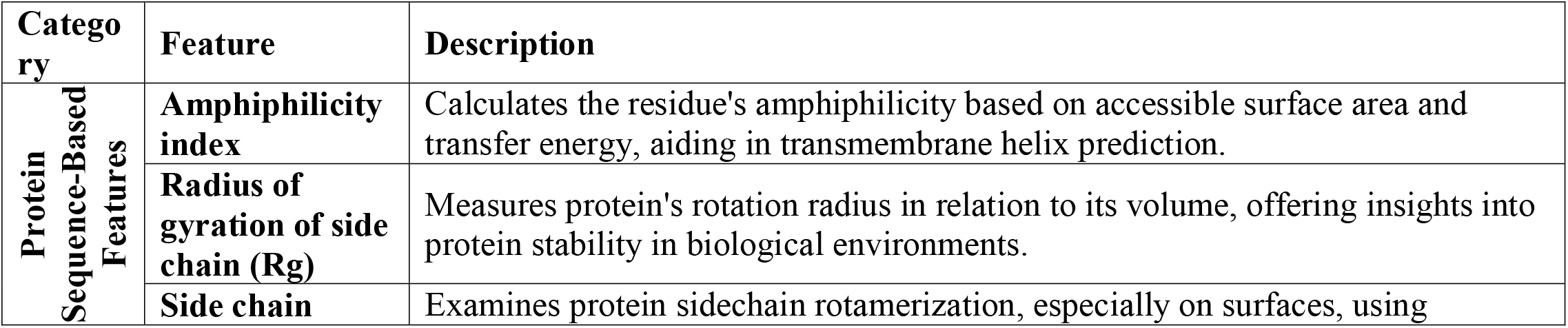
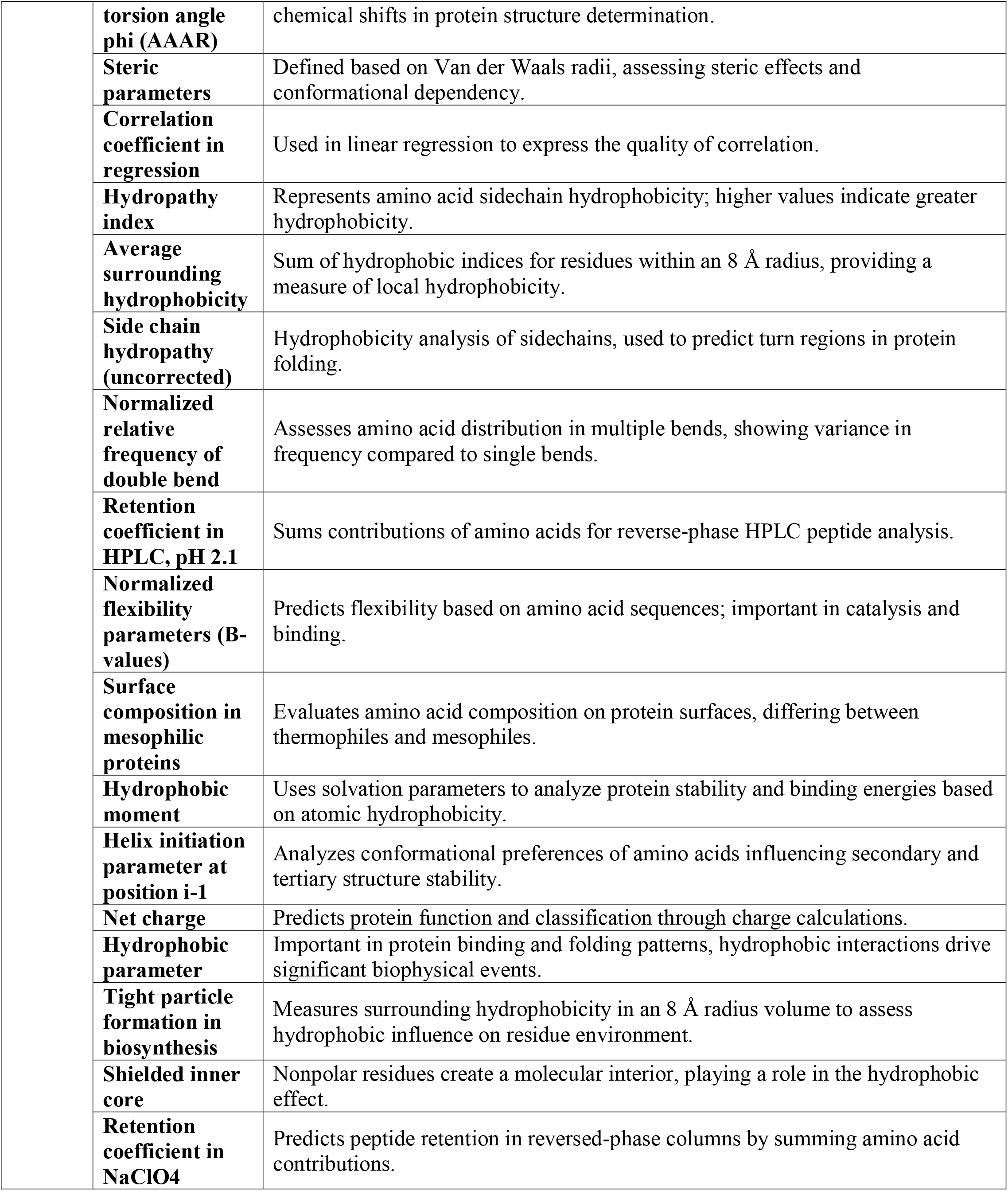
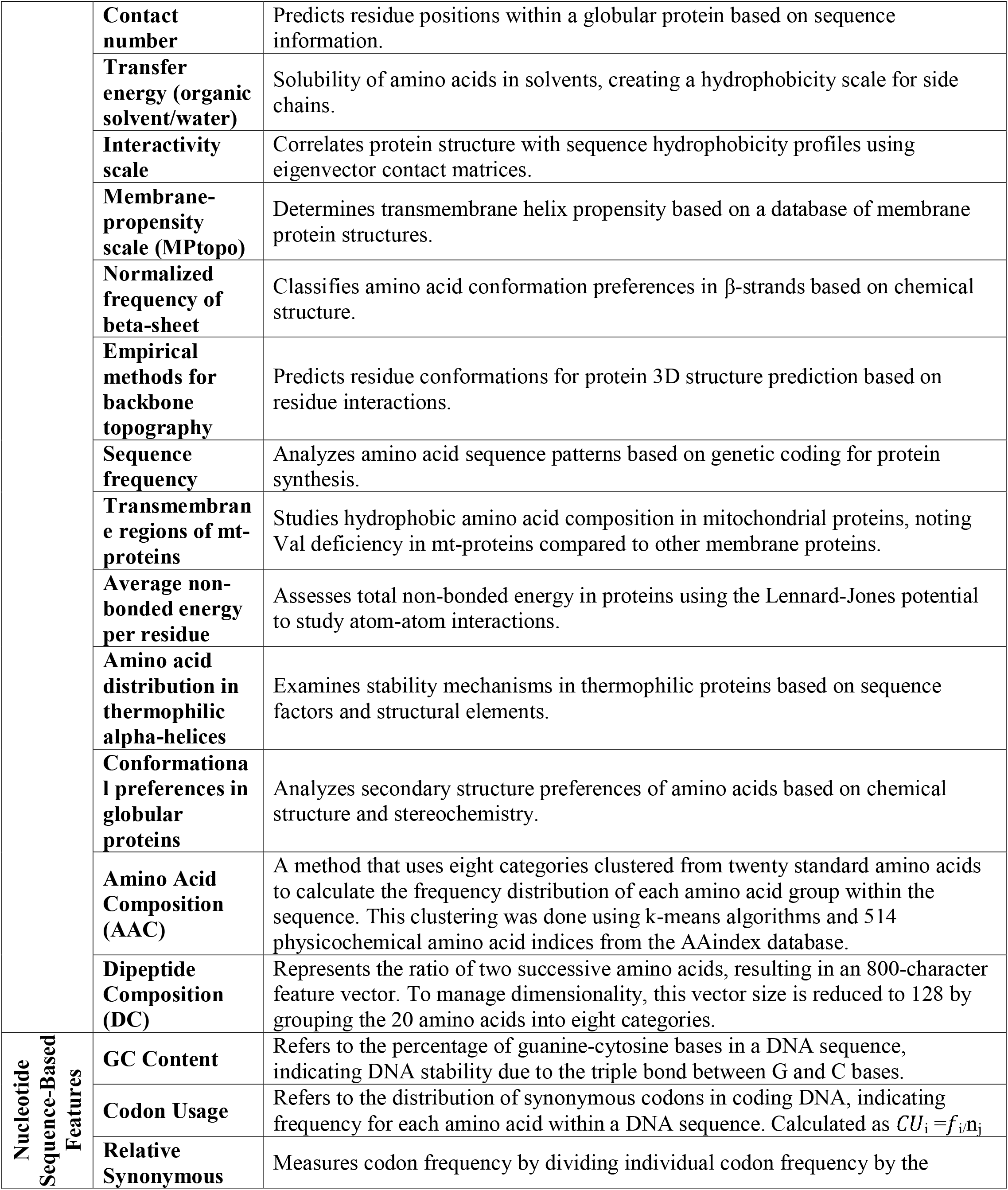
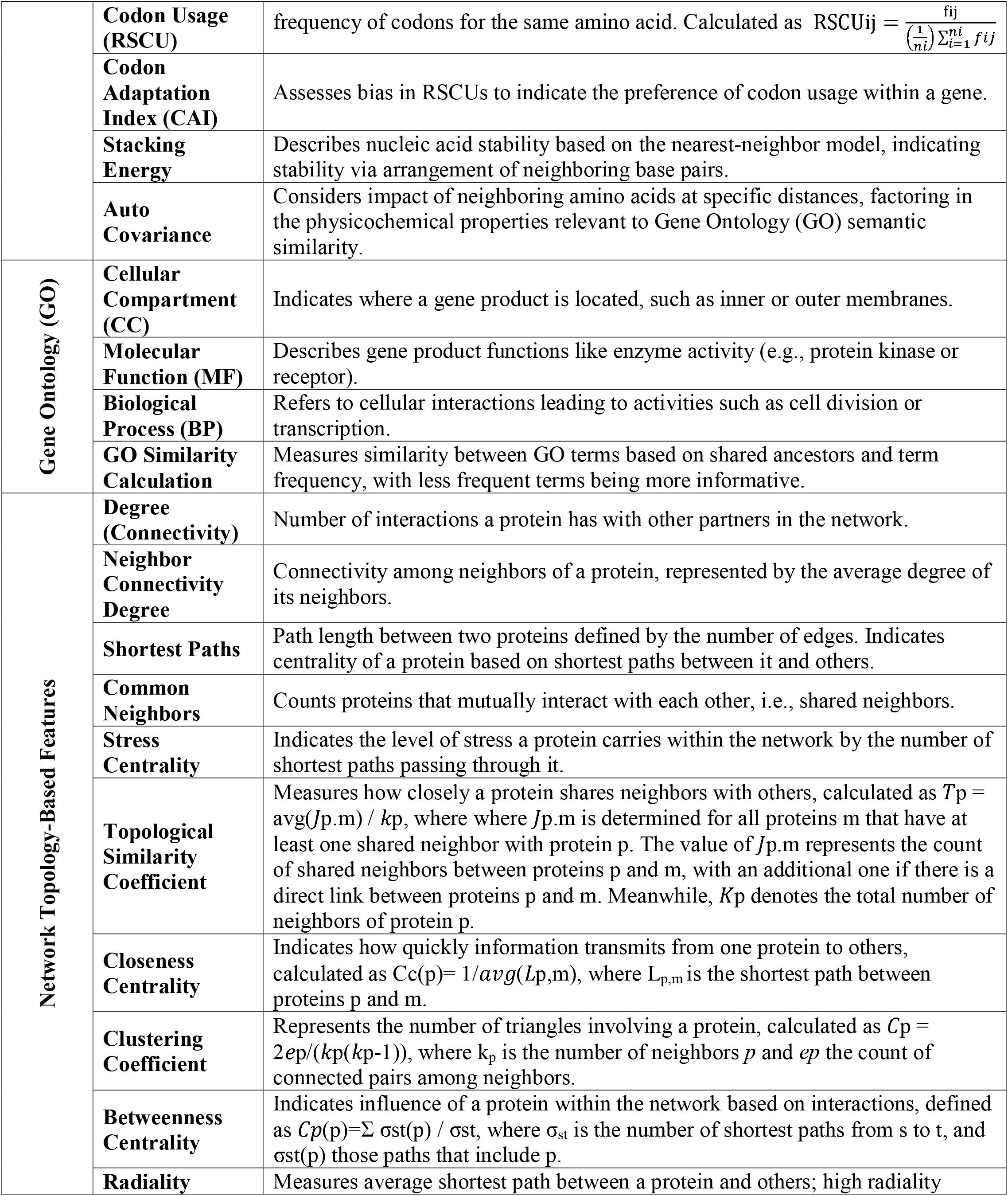
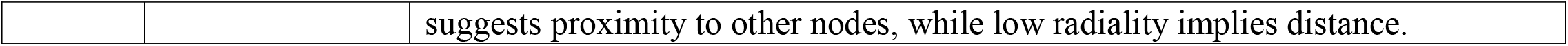
Description of features used in this study

### Predictive Model Development and Evaluation

Five machine learning classifiers were utilized: Decision Tree (DT), Support Vector Machine (SVM), Naive Bayes (NB), K-Nearest Neighbor (KNN), Logistic Regression (LR), and Random Forest (RF). The performance of each model was evaluated using metrics including accuracy, sensitivity, specificity, positive predictive value (PPV), negative predictive value (NPV), and Prevalence Rate. Additionally, we applied stratified k-fold cross-validation (k = 10) to assess the generalizability of the predictive models. The stratification ensured that the ratio of positive to negative samples was preserved in each fold.

## Results

Retroviruses interact with host proteins through a range of mechanisms that allow them to manipulate cellular entry, transport, DNA integration, and protein synthesis. These interactions enable the virus to establish a persistent infection and highlight the complex interplay between virus and host that is essential to retroviral replication and survival.

In this study, we identified key viral and human protein hubs within the virus-host interaction network. The viral protein with the highest number of interactions were ENV_HV1H2 (275 interactions), REV_HV1BR (203 interactions), and TAT_HV1H2 (132 interactions), highlighting their central roles in viral pathogenesis. For human proteins, CD4 and HCK (9 interactions each) along with CCNT1 and DDB1 (6 interactions each) emerged as the primary interaction hubs, indicating their potential significance in retroviral infection.

Our machine learning models - Logistic Regression (LR), Support Vector Machine (SVM), Naive Bayes (NB), Decision Tree (DT), and Random Forest (RF)-were evaluated across metrics including accuracy, sensitivity, specificity, positive predictive value (PPV), negative predictive value (NPV), and prevalence rate. Among the models, DT achieved the highest sensitivity (77%), while LR and KNN had the highest specificity (52% and 51%, respectively). Accuracy ranged from 56% for NB to 61% for LR and KNN, highlighting these models’ overall robustness. In terms of PPV and NPV, LR and KNN again demonstrated superior performance, with values of 63% and 57%, respectively. Prevalence rate values further confirmed LR and KNN as the top models, both scoring 63% (Figure 1). These results suggest LR and KNN as viable models for reliable retroviral interaction prediction.

**Figure 1.**
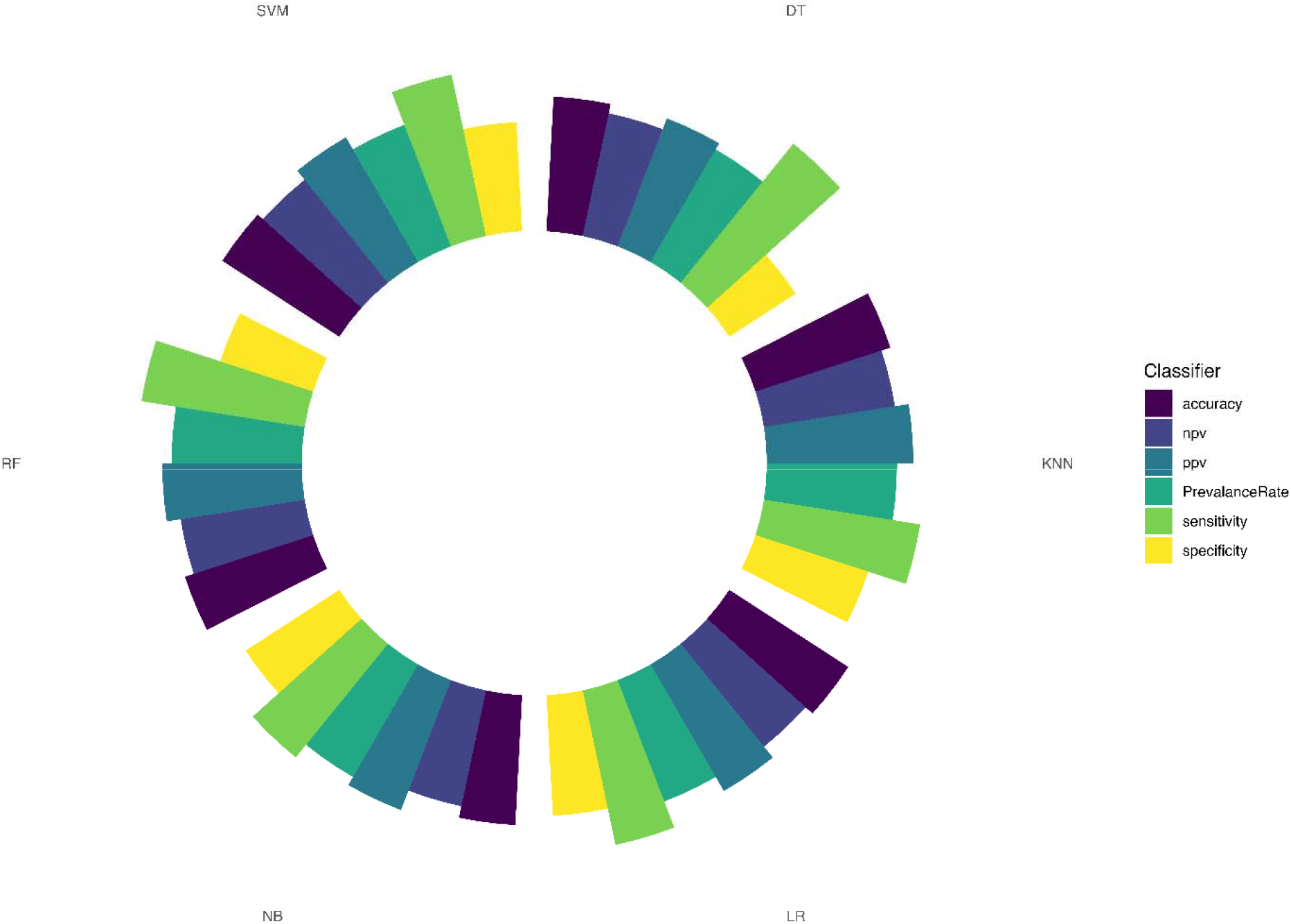
Results of various models (LR, SVM, NB, DT and KNN) that were evaluated by sensitivity, specificity, PPV, NPV, AUC and Prevalence Rate)

Feature importance analysis, displayed in Figure 2, highlighted GC content, Gene Ontology semantic similarity, hydropathy index, and net charge as the most influential predictors of retroviral interactions.

**Figure 2.**
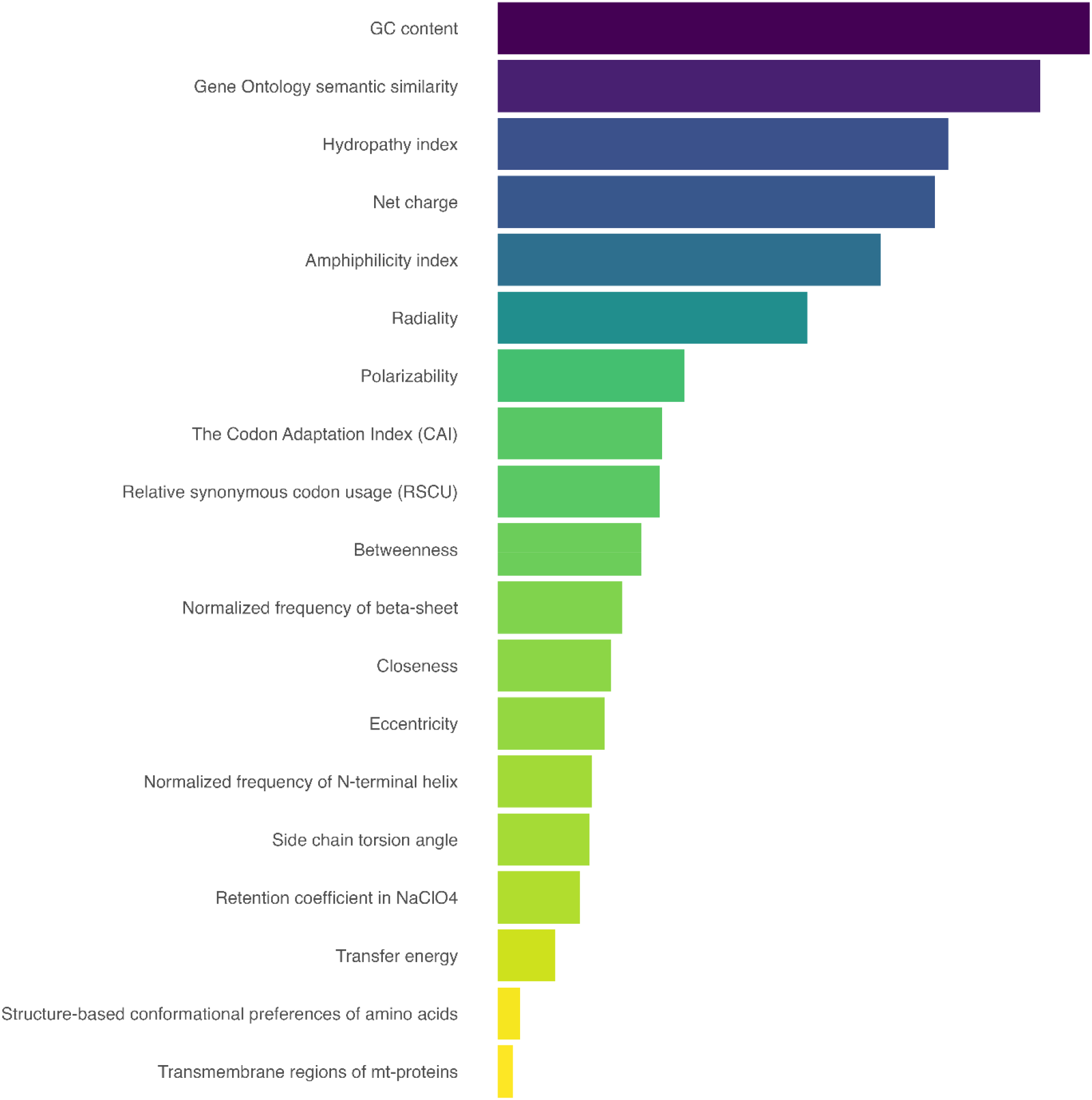
All features were sorted from most main feature to lowest main feature and ten top were selected for next studies

After conducting the feature importance analysis, we trained the models using the key features and evaluated their performance metrics. The radar chart provides a comparison between models trained on both key and full feature sets. The results indicated that using important features led to more accurate predictions across most classifiers, without a loss of efficiency when compared to models trained on all features (Figure 3).

**Figure 3.**
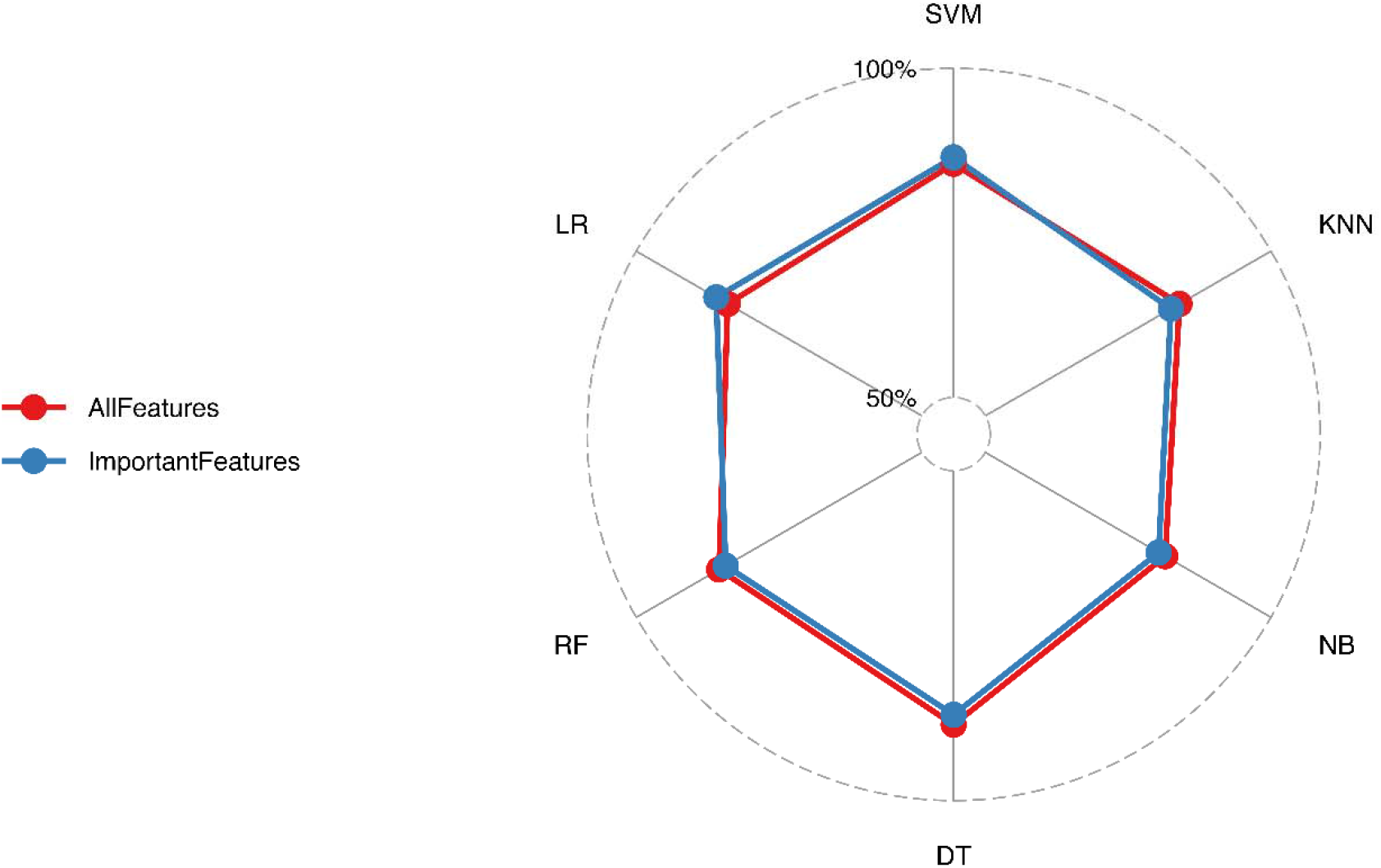
Radar chart illustrating the comparative performance of LR, SVM, KNN, NB, RF, and DT classifiers when trained on important features versus all features.

## Discussion

Numerous ML studies have been conducted to predict the VHPPI, employing various models and strategies aiming to improve prediction accuracy. These models differ in their negative dataset, feature, and feature selection.

In terms of dataset construction, there are few studies where experimentally approved data for both negative and positive datasets. For instance, in a study conducted to predict human-HCV protein interactions, an ensemble model combining RF, NB, SVM, and multilayer perception (MLP) classifiers were employed. The prediction was done using amino acid composition, post-translational modifications, and network centrality measures as biologically relevant features. The study gathered experimentally approved non-interacting proteins which avoid false negatives. This careful negative sampling, combined with the ensemble approach, yielded high specificity (92%) and sensitivity (84%) [41].

Meanwhile, since experimentally validated non-interacting virus-human PPIs are relatively rare, most studies employed pseudo-negative sampling. For instance, Barman et al. predicted the VHPPI for hepatitis virus using an RF-based model trained by features of domain-domain associations and amino acid composition. Their negative dataset was constructed by random sampling of non-interacting protein pairs. This method carries the risk of increasing false negatives. However, the RF model demonstrated reasonable predictive powers (71.69%) [42]. Similarly, Tsukiyama et al. employed non-overlapping pairs as pseudo-negative samples for their deep learning Long Short-Term Memory (LSTM) model which achieved very high accuracy (98.4%) [43].

Considering the significant important of several viruses, some researchers preferred to computationally study PPI for a strain, a family, or a genus. For instance, Khorsand et al. studied SARS-CoV-2-human protein-protein interaction network during the pandemic [44]. Moreover, a Siamese neural networks study has conducted to predict the virus host PPI for SARS-CoV-2 and JCV [45].

## Conclusion

This study demonstrates the effectiveness of computational approaches in predicting virus-human protein-protein interactions, with retroviral PPIs as a case study. Using carefully constructed positive and pseudo-negative datasets, we identified and evaluated key protein features critical for interaction prediction. Among the machine learning models tested, Logistic Regression and K-Nearest Neighbor achieved the most promising results in terms of sensitivity and specificity, suggesting their utility in retroviral interaction prediction. Furthermore, our feature importance analysis underscores the value of biologically relevant features, which can improve model accuracy while reducing computational costs. These findings pave the way for refined predictive models that can be applied to various viral families, enhancing our ability to understand viral pathogenesis and identify therapeutic targets.

